# Prokaryotic Gabija complex senses and executes nucleotide depletion and DNA cleavage for antiviral defense

**DOI:** 10.1101/2023.05.02.539174

**Authors:** Rui Cheng, Fengtao Huang, Xueling Lu, Yan Yan, Bingbing Yu, Xionglue Wang, Bin Zhu

## Abstract

The Gabija antiviral system consists of the GajA and GajB proteins. We previously revealed that GajA is a DNA nicking endonuclease. In this work, we found that the DNA binding of GajA is strictly inhibited by NTP. Furthermore, the antiviral defense of GajA requires the assistance from GajB, which senses DNA termini produced from the DNA nicking by GajA to hydrolyze (d)A/(d)GTP. The synergy between the DNA cleavage by GajA and the nucleotide hydrolysis by GajB results in an efficient abortive infection defense against virulent bacteriophages. GajA binds to GajB to form stable complexes *in vivo* and *in vitro*. However, a functional Gabija complex requires the molecular ratio between GajB and GajA below 1:1. Through (i) sequential sensing and executing the nucleotide depletion and DNA cleavage to cause a cascade suicide effect and (ii) stoichiometry regulation of the DNA/nucleotide processing complex, the Gabija system exhibits a unique mechanism distinct from other known prokaryotic antiviral systems.

## INTRODUCTION

To resist frequent and varied attacks by viruses, prokaryotes have evolved numerous ingenious defense strategies that are collectively known as the prokaryotic “immune system” (1–3). The most widespread prokaryotic immune systems are the adaptive immune system CRISPR/Cas, which provides acquired immunity through memorization of past phage attacks (4), and innate restriction-modification (R-M) immune systems that target specific, predefined DNA sequences of invading phages (5). Gene editing technology originated from CRISPR/Cas systems has been very influential. Similarly, restriction endonucleases derived from R-M systems have revolutionized recombinant DNA technology (6).

The recent boom in metagenomic analyses has implied that a diverse array of unknown defense systems exist in bacteria (7–16). To date, many novel prokaryotic antiviral mechanisms have been revealed, including but not limited to cyclic oligonucleotide-based antiphage signaling system (CBASS) (17–21), bacteriophage exclusion (BREX) system (22,23), prokaryotic Argonautes (pAgos) (24), defence island system associated with restriction-modification (DISARM) (25), nuclease-helicase immunity (Nhi) (26), and SspABCD–SspE system (27). Many of these newfound systems defend against phage infection via an abortive infection mechanism, e.g., phage anti-restriction-induced system (PARIS) (28), pyrimidine cyclase system for antiphage resistance (Pycsar) (29), Helicase-nuclease Abi (hna) (30), DarTG toxin-antitoxin system (31), and DdmABC and DdmDE (32). The abortive infection systems protect the bacterial population but not individual cells because they cause cell suicide before phage progeny release (33–37). These investigations considerably expanded our knowledge of bacterial immunity.

Doron et al. identified 10 novel prokaryotic antiviral systems (8). Except for the recently characterized Thoeris system (38,39) and Wadjet system (40,41), the antiviral mechanisms of these systems remain elusive. Among these systems, we focus on the Gabija defense system, which comprises two predicted genes, GajA and GajB. The Gabija system from *Bacillus cereus* VD045 confers resistance to a broad range of phages, including phages phi29, rho14, phi105, and SpBeta (8). Bioinformatics analysis suggested that Gabija is widespread in prokaryotes, existing in 4,360 sequenced genomes, about 8.5% of all analyzed genomes (8). By contrast, CRISPR/Cas systems are found in about 40% of all of the sequenced bacteria (42,43), and R-M systems are found in about 75% of prokaryote genomes (44).

Many known bacterial defense systems attack bacteriophage genomic DNA and most of their elements, such as the CRISPR/Cas and R-M systems, have the ability to specifically process nucleic acids (4,5). Our recent study has revealed that one component of the Gabija system, GajA is a nucleotide-sensing and sequence-specific DNA nicking endonuclease (45). However, the role of the other component, GajB, which is predicted to be a UvrD-like helicase, remained unclear (8). As previously reported, UvrD can function either as a helicase or only as a single-stranded DNA (ssDNA) translocase, which translocates in a 3ʹ to 5ʹ direction using its ssDNA-dependent ATPase activity (46–48). 5ʹ-ss-duplex DNA junctions can serve as loading sites for the monomeric UvrD translocase (49). Recent research further implied that UvrD facilitates DNA repair by pulling RNA polymerase backward (50).

In this work, we carried out extensive genetic and biochemical investigations on the Gabija defense system and reported its molecular mechanism. We found that the Gabija system exhibits broad and efficient phage resistance through an abortive infection manner and both GajA and GajB are essential for its function. GajA exhibited specific oligomerization and DNA binding properties, the latter being strongly inhibited by NTP. Unexpectedly, GajB exhibits strong (d)A/(d)GTP hydrolysis activity but no helicase activity. We identified the substrates and optimal reaction conditions of GajB and further revealed that GajB activity is activated by DNA termini. GajA and GajB form a stable complex, in which the DNA nicking by GajA stimulates GajB activity, and at a certain molecular ratio, GajB enhances the DNA binding and cleavage of GajA. Our results demonstrate a novel mechanism in which a nucleotide-sensing DNA nickase and a DNA-termini-dependent (d)A/(d)GTPase coordinate to mediate antiviral defense.

## RESULTS

### The Gabija antiviral defense requires both GajA and GajB and functions through abortive infection

The Gabija defense system was initially predicted to consist of two components, GajA and GajB (Figure 1A) (8). To confirm its antiviral function, we cloned the Gabija gene cassette sequence (GajAB coding sequence, from the start codon of GajA gene to the stop codon of GajB gene, located at 94190–97412 of the *B. cereus* VD045 genome [AHET01000033]), the GajA gene alone, or the GajB gene alone, each under the control of an *E. coli* T5 promoter, into the host strain *E. coli* B (ATCC^®^ 11303™), which naturally lacks the Gabija system. We then challenged the transformed strain with three coliphages and found that the system conferred protection against all three typical types of lytic phages: T7 (Podoviridae), T4 (Myoviridae), and T5 (Siphoviridae) (Figure 1B). The Gabija system exhibited broad phage resistance, showing very strong resistance to phages T7 and T4 (>10^7^ fold decrease in efficiency of plating [EOP]) and weaker resistance to phage T5 (∼10^3^ fold decrease in EOP) (Figure 1B). In a previous report, the two-gene system together with its flanking intergenic regions (located at 93871–97763 of the *B. cereus* VD045 genome [AHET01000033], Figure 1A) showed defense against phages (8). Our data further confirmed that only the coding region is essential for defense against phages. Both GajA and GajB genes in the system appear to be essential for its functionality, because deletion of either gene resulted in the loss of protection from phage infection (Figure 1B).

**Figure 1.**
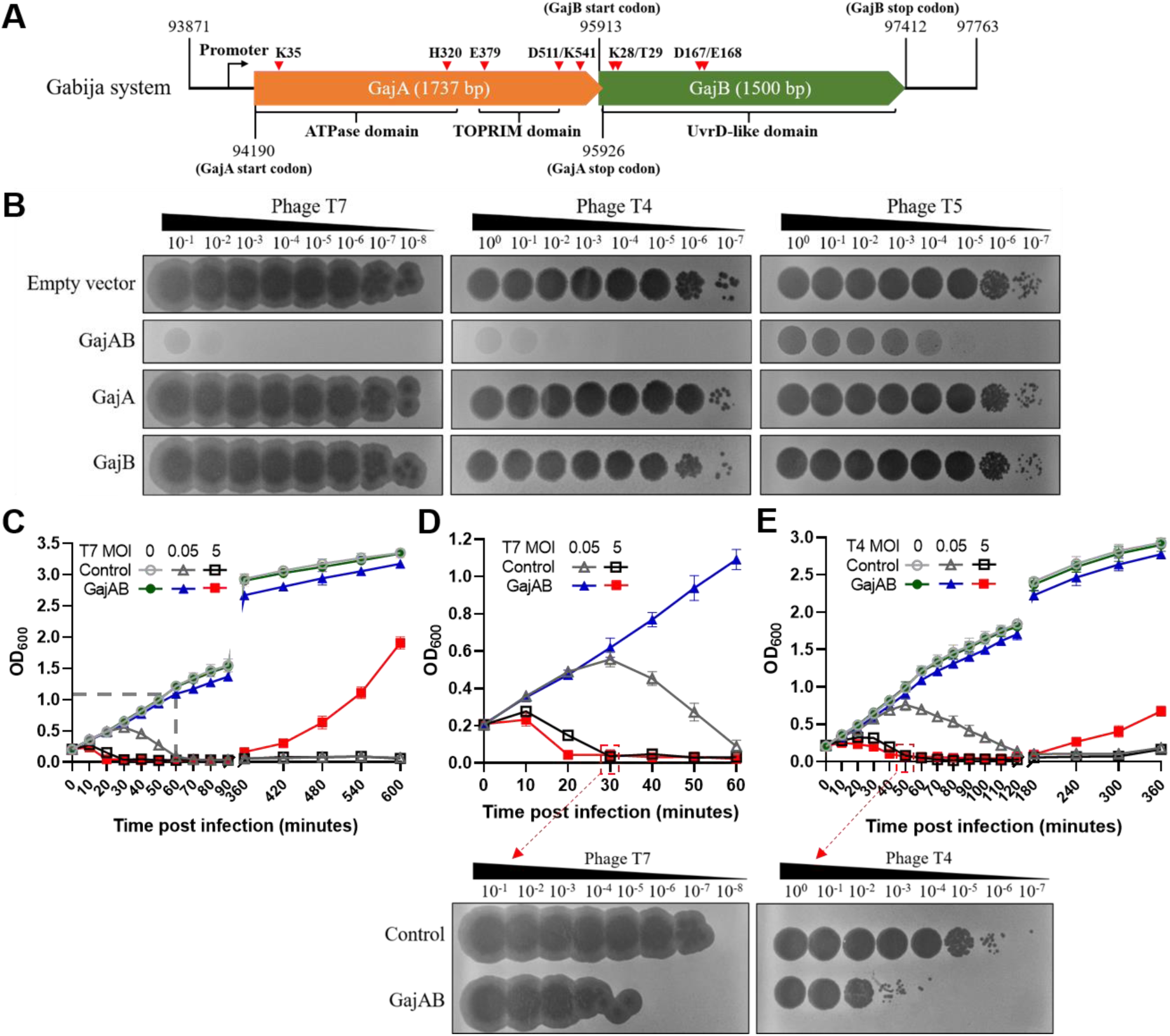
The Gabija bacterial defense system from *B. cereus* VD045 has cross-species functions in *E. coli* to abolish phage infection. (**A**) The *B. cereus* VD045 Gabija gene cassette contains two predicted genes, GajA (1737 bp) and GajB (1500 bp) with a 14-bp overlapping. The range of the Gabija gene cassette in the *B. cereus* VD045 genome [AHET01000033], the promoter upstream of GajA gene, positions of the start codon and stop codon for GajA and GajB genes, the predicted domains of GajA and GajB, as well as positions of the key residues investigated by mutagenesis in this work, are labeled in the schematic. (**B**) Phage infection of *E. coli* B containing pQE82L plasmids inserted with various Gabija genes and a control plasmid (empty vector). Shown are 10-fold serial dilutions of plaque assays with phages T7, T4, and T5. The sequences of the Gabija gene cassette including genes GajA and GajB (located at 94190**–**97412 of the *B. cereus* VD045 genome [AHET01000033], GajAB), the GajA gene alone, or the GajB gene alone, as shown in (**A**). The recombinant vectors were transformed into *E. coli* B. Empty vector was used as a negative control. Plates were spotted with 4 μl of the three phages diluted in LB at eight 10-fold dilutions (10^−1^–10^−8^ for T7 and 10^0^–10^−7^ for T4 and T5) and incubated at 37°C overnight before images were taken. Images are representative of three replicates. (**C–E**) Growth curves of Gabija-expressing (solid) and control (empty) cultures with and without infection by phages T7 (**C**) and T4 (**E**) at an MOI of 0, 0.05, or 5. Each line represents the mean ± SD of three independent experiments. OD_600_, optical density at 600 nm. (**D**) Magnification of the area marked with a dashed rectangle in **C** (MOI of 0.05 and 5). To detect the release of phage progeny by dead cells with or without the Gabija defense system, the lysates of dead cells with phage infection at MOI 5 (30 min post-infection for T7 and 50 min post-infection for T4) from the above assays were collected. Then 10-fold serial dilutions of the lysates (4 μl) were spotted on *E. coli* B carrying empty vector to compare their plaque efficiency. Plates were incubated at 37°C overnight before images were taken. Images are representative of three replicates.

Many bacterial defense systems are known to protect bacteria by triggering cell death to terminate phage infection, which is known as abortive infection (33). Our previous investigation revealed that GajA efficiently catalyzes DNA nicking on both T7 and host genomic DNA *in vitro* (45), indicating that Gabija may be an abortive infection system. To clarify the defense mode of the Gabija system, phage T7 or T4 was mixed with *E. coli* B harboring the empty vector (control) or the Gabija system (GajAB) in liquid media at an MOI of 0, 0.05, and 5 and cell growth was tracked by measuring OD_600_. Gabija-mediated protection against phage T7 or T4 infection was clearly observed at an MOI of 0.05 (Figure 1C–E). However, at an MOI of 5, the collapse of the Gabija-containing cells occurred earlier than the phage-induced lysis observed in Gabija-lacking cells (Figure 1C–E). Moreover, Gabija-expressing cells were able to recover several hours after the collapse, in contrast to control cells, which did not recover following phage lysis at the same time points (Figure 1C–E). Furthermore, we collected lysates of the dead cells with and without the Gabija system in the MOI 5 groups at 30 min (for T7) or 50 min post-infection (for T4) from the above assays and compared their plaque-forming efficiency (Figure 1D, E, bottom). All the lysates formed plaques, likely due to unabsorbed phages from the initial infection at MOI 5. However, the plaque-forming efficiency of the lysates from dead cells with the Gabija defense was 100-fold (T7) or 1000-fold (T4) lower than that from dead cells without the Gabija defense (Figure 1D, E, bottom), indicating that phage progeny was not released by the dying cells containing the Gabija system. These data confirmed that Gabija defends through abortive infection, leading to the death of infected cells before the maturation of phage progeny.

We examined the role of conserved residues of GajA and GajB in antiviral defense. In previous work, we characterized that GajA is a nucleotide-sensing DNA nicking endonuclease (Figure 2A), utilizing a TOPRIM domain to cleave DNA and an ATPase-like domain to mediate the regulation by nucleotide concentration (45). Therefore, we generated three mutations, E379A, D511A, and K541A, in the TOPRIM domain and two mutations, K35A and H320A, and a truncation (GajA-CTR, 1–347 amino acid deletion) in the ATPase-like domain, introduced these mutations into the GajAB gene cassette, and tested their effects on phage defense. We found that all of the above mutations abolished the antiviral function of the Gabija system (Figure S1). E379A, D511A, and K541A have been shown to inactivate the DNA cleavage of GajA and H320A affects the NTP inhibition of GajA (45); however, the K35A mutation, without any observed effect on the DNA cleavage activity of GajA, also abolished the defense function of the whole system. To target conserved amino acid residues of GajB, sequence alignment was performed between GajB and known helicases DrUvrD, BsPcrA, EcUvrD, and EcRep (47, 51) (Figure S2). Double mutations of the key residues of UvrD helicases in GajB, K28A/T29A or D167A/E168A, resulted in the loss of phage resistance (Figure S3). These results suggest that specific activities of both GajA and GajB are essential for the Gabija defense system.

**Figure 2.**
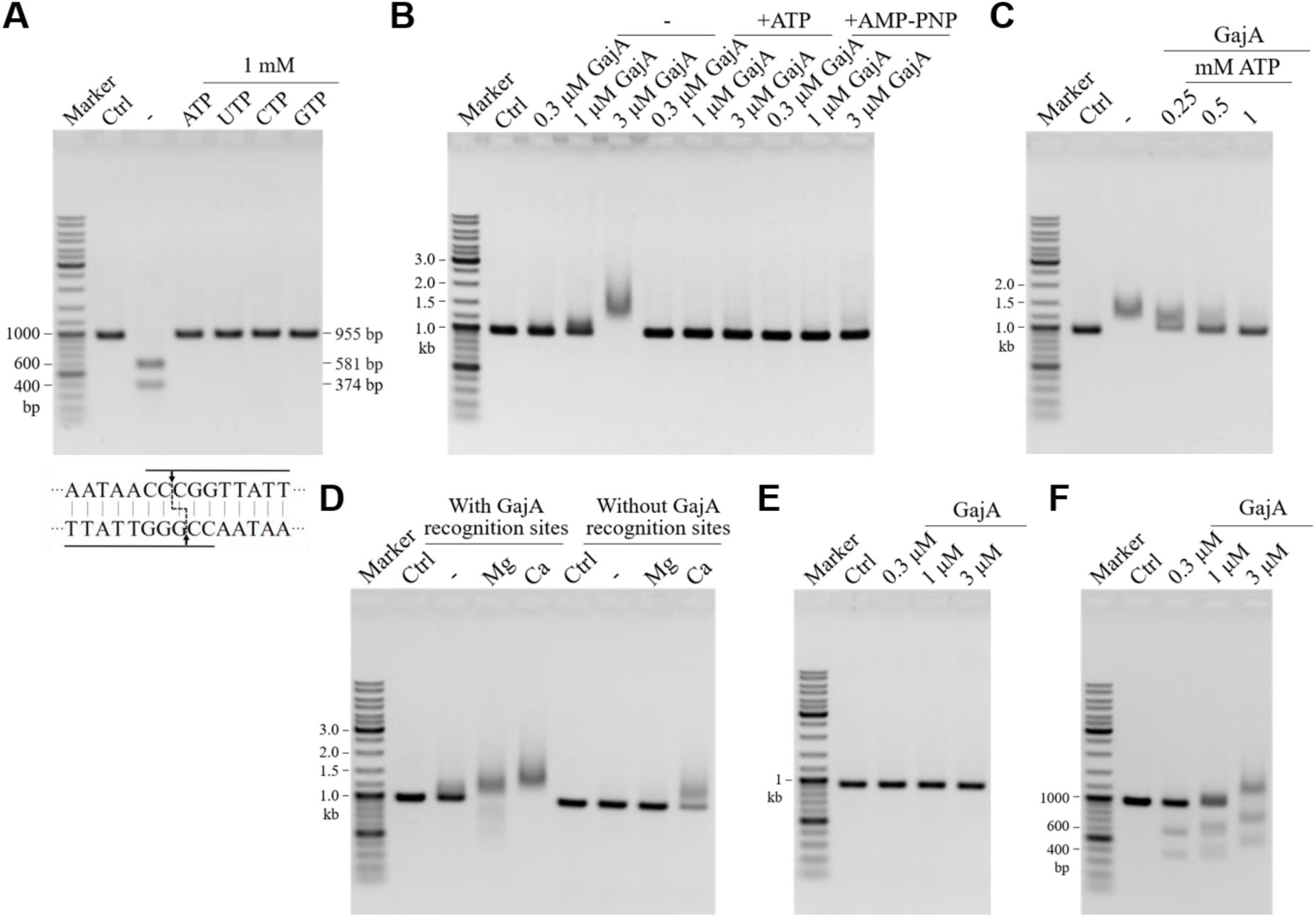
The DNA binding of GajA as determined by EMSA. (**A**) Effects of NTP on GajA endonuclease activity. The DNA substrates pUC19*-*955 were amplified from plasmid pUC19 by primers pUC19-F/R (listed in Table S2) containing two overlapping GajA recognition sequences *AATAA**CCCGG*****TTATT** (marked by lines). Nicking by GajA closely on both strands results in the cleavage of the DNA into two pieces (581 bp and 374 bp). 125 ng of pUC19*-*955 DNA was incubated with 0.2 µM GajA in a final volume of 10 µl in the optimal reaction buffer. Reactions were performed at 37°C for 5 min and then stopped by the addition of 2 µl of 6× loading dye containing 20 mM EDTA. “-” indicates the reaction without NTP. The cleavage patterns are shown on the bottom of the gel. Each of the four NTPs (1 mM) was added to the reactions. (**B**) The DNA binding by GajA (0.3, 1, and 3 μM) in the absence or presence of 1 mM ATP or AMP-PNP in binding buffer containing 5 mM CaCl_2_. Lanes labeled with dashes indicate no ATP or AMP-PNP addition. (**C**) Inhibition of the DNA binding of GajA by ATP. Reactions were performed with 3 µM GajA in the absence or presence of various amounts of ATP in binding buffer containing 5 mM CaCl_2_. (**D**) The DNA binding of GajA in the presence or absence of Mg^2+^ or Ca^2+^. Reactions contained 3 µM GajA. For **B**–**D**, 125 ng of pUC19*-*955 DNA was used as the substrate. (**E, F**) The binding to and cleavage of two DNA substrates without (**E**) or with (**F**) GajA recognition sites (*AATAA**CCCGG*****TTATT**) by GajA (0.3, 1, and 3 μM) in binding buffer containing 5 mM MgCl_2_. For **B**–**F**, reactions were incubated at 4°C for 30 min before adding 2 µl of 6× loading dye containing 20 mM EDTA. For **A**–**F**, Ctrl refers to the control without protein. Samples were analyzed via native agarose gel electrophoresis and ethidium bromide staining.

### ATP regulates DNA cleavage activity of GajA by inhibiting its specific DNA binding

Our previous study demonstrated the inhibition of DNA cleavage activity of GajA by NTP (Figure 2A) (45), but the underlying mechanism was unclear. In this work, we detected the DNA binding of GajA in the absence or presence of ATP or AMP-PNP. On the pUC19-955 substrate containing GajA recognition sites, GajA showed DNA cleavage activity in the presence of Mg^2+^ (Figure 2A) but not in the presence of Ca^2+^ (45). Thus we used Ca^2+^ instead of Mg^2+^ (45) in the reactions to detect the DNA binding of GajA without substrate cleavage. The electrophoretic mobility shift assay (EMSA) demonstrated the binding of GajA to a DNA substrate containing two overlapped GajA recognition sites *AATAA**CCCGG*****TTATT** (one recognition site in the plus strand shown in bold and the other in the minus strand shown in italics), which was inhibited by ATP and AMP-PNP (Figure 2B). The DNA binding of GajA was sensitive to the ATP concentration and completely inhibited by 1 mM ATP (Figure 2C). Ca^2+^ supported a specific DNA binding of GajA in the presence of GajA recognition sites and also supported a weaker non-specific DNA binding of the GajA in the absence of recognition sites (Figure 2D). Mg^2+^ only supported the specific DNA binding of GajA (Figure 2D), showing no binding on a DNA substrate amplified from the pUC19 plasmid containing no GajA recognition sequence (Figure 2E). When the substrates containing two overlapped recognition sites were cut into two fragments in the presence of Mg^2+^, GajA was still bound to the cleaved DNA fragments (Figure 2F), which is consistent with a previously reported inefficient turnover of GajA (45). The GajA-bound DNA appears to be a clear gel band shift rather than a smear, suggesting a specific stoichiometry of GajA/DNA in the complex.

Subsequently, we detected the DNA binding activity of GajA and its mutants in the absence or presence of ATP. K35A, H320A, and D511A mutations decreased the inhibition of GajA DNA binding activity by ATP. K541A mutation enhanced the DNA binding of GajA and changed the gel shift of the GajA/DNA complex, indicating that the mutation altered the binding mode or the stoichiometry in the complex. Like that for the WT, ATP inhibited the DNA binding of all of the GajA mutants, except for E379A (Figure S4A). GajA-E379A was peculiar in that its DNA binding resulted in a wider range of gel shifts and was not inhibited by ATP (Figure S4A). Further, we found that the DNA binding of GajA-E379A was not dependent on recognition sites and metal ions (Figure S4B). The non-specific DNA binding of GajA-E379A indicated that the E379 site is vital to determine the specificity of GajA for recognition sites.

### GajB forms a complex with GajA *in vitro* and *in vivo*

To investigate the relationship between GajA and GajB, we constructed the predicted coding sequence (AHET01000033.1: 94,190–97,412, sequence listed in Table S1) of GajAB (WT Gabija gene cassette). In this construction, GajA was fused to an N-terminal 6×His-tag. The recombinant proteins were purified on a Ni-NTA agarose column. Interestingly, non-tagged GajB was co-purified along with His-tagged GajA (Figure S5A). We also constructed another vector GajA+B (His-tagged GajA and non-tagged GajB overexpressed under the control of two individual promoters). In this case, GajB was also co-purified with GajA (Figure 3A and Figure S5B). To verify the *in vivo* results, we performed *in vitro* assembly of the GajA/B complex. The predicted coding sequence for GajA or GajB alone was cloned into pET28a vectors and individual protein was overexpressed, respectively. The supernatants containing His-tagged GajA and non-tagged GajB were mixed before loading onto a Ni-NTA agarose column. GajB was again co-purified with GajA (Figure S5C). Collectively, these data demonstrated a stable binding between GajB and GajA.

**Figure 3.**
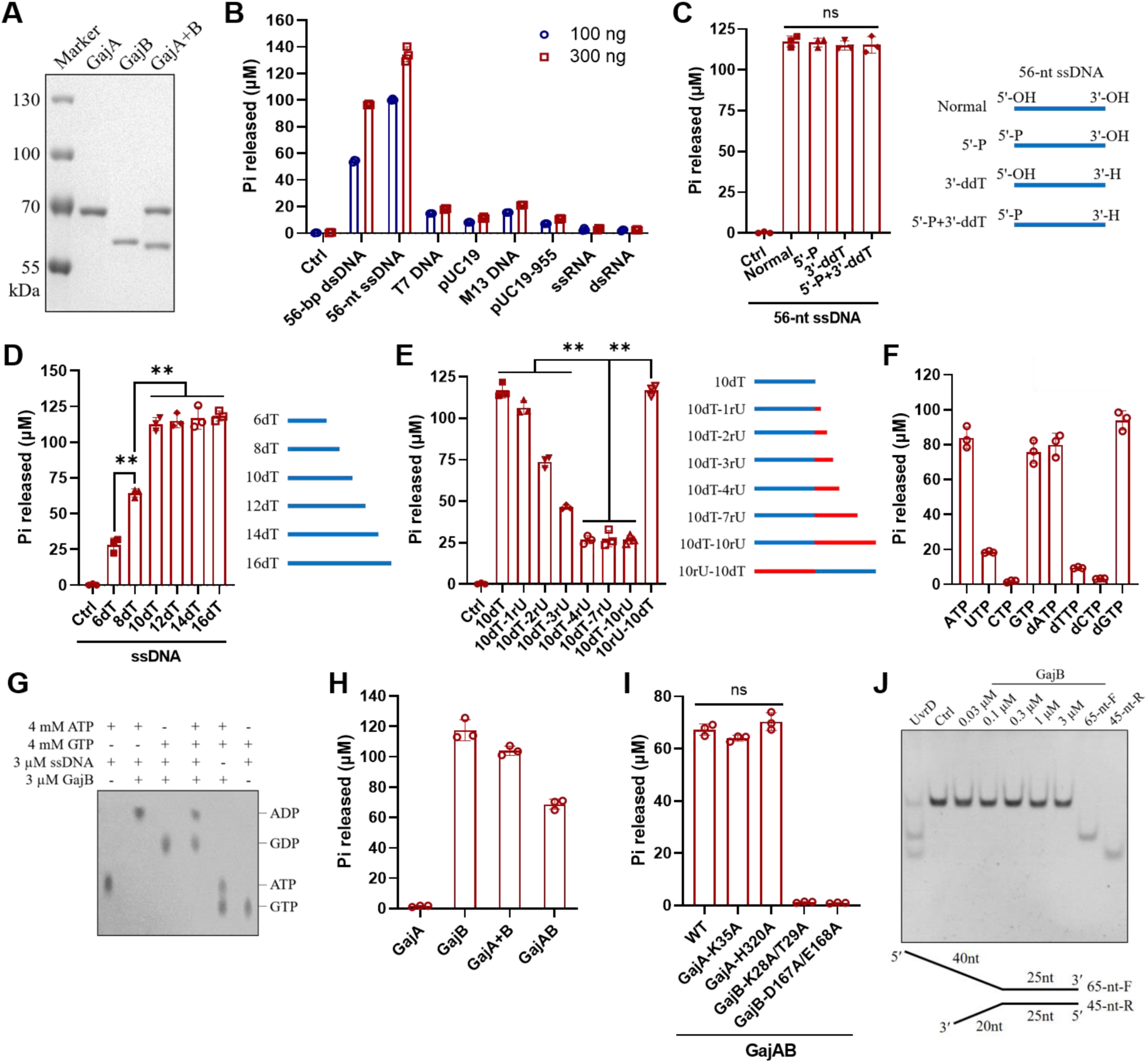
DNA terminus-dependent (d)A/(d)GTPase activity of GajB. (**A**) SDS-PAGE gel showing purified GajA (with an N-terminal His-tag), GajB (with an N-terminal His-tag), and GajA+B (His-tagged GajA and co-purified non-tagged GajB). (**B**) ATP hydrolysis by GajB in the presence of various DNA and RNA, including 56-bp dsDNA, 56-nt ssDNA, T7 DNA, pUC19 plasmid DNA, M13 DNA, PCR-amplified pUC19-955 DNA, 600-nt ssRNA, and 600-bp dsRNA (sequences are listed in Table S3). 56-bp dsDNA was prepared by annealing two complementary 56-nt ssDNA (56-nt ssDNA-F and 56-nt ssDNA-R). Similarly, 600-bp dsRNA was prepared by annealing two complementary 600-nt ssRNA. Reaction mixtures containing 20 mM Tris-HCl pH 7, 1 mM DTT, 10 mM MgCl_2_, 0.5 mM ATP, 0.5 μM GajB, and 100 ng or 300 ng DNA or RNA were incubated at 30°C for 15 min. (**C**–**E**) ATP hydrolysis by GajB in the presence of various ssDNAs: (**C**) 56-nt ssDNA with various terminal groups; (**D**) poly-dT ssDNA of various lengths; and (**E**) 10-nt poly-dT ssDNA with various rU attachments. Schematics of various ssDNAs are shown on the right of each panel. Reaction conditions were similar to those in (**B**) except for the nucleic acids (0.5 μM) added as indicated in the graph. For **B**–**E**, reactions without DNA/RNA added (Ctrl) were included as a control. (**F**) Hydrolysis of various NTPs and dNTPs by GajB. Reaction mixtures containing 20 mM Tris-HCl pH 7, 1 mM DTT, 10 mM MgCl_2_, 0.5 μM 56-nt ssDNA, 0.5 μM GajB, and 0.5 mM of each NTP or dNTP were incubated at 30°C for 15 min. (**G**) TLC analysis of the A/GTPase products of GajB. Reaction conditions were similar to those in (**F**). GajB effectively hydrolyzes ATP/GTP to ADP/GDP and Pi. (**H**) ATPase activity of various combinations of Gabija proteins as indicated at the bottom of the graph. All reactions containing 0.5 mM ATP, 0.5 μM 56-nt ssDNA, and 0.5 μM total protein in optimal reaction buffer were incubated at 30°C for 15 min . (**I**) ATPase activity of GajAB and its mutants. Reaction conditions were similar as in (**F**) except that various proteins as indicated at the bottom of the graph were tested. (**J**) GajB displayed no DNA unwinding activity. The Y-type dsDNA substrate was used as helicase substrate, which was prepared by annealing two oligonucleotides (sequences are listed in Table S2). Reactions were performed as described in the Materials and Methods. UvrD was used as a positive control. Pi release was monitored by the PiColorLock™ kit. Bar graphs represent the average of three independent experiments with error bars representing the standard error of the mean. Statistical significance was calculated using Student’s *t*-test, which is indicated as follows: ns, not statistically significant, \*\**P* < 0.005.

### GajB senses DNA 3ʹ termini for ATP hydrolysis

Bioinformatics analysis predicted that GajB is a UvrD-like helicase. To elucidate its function, GajB (>90% homogeneity) was purified as an N-terminal His-tagged protein (Figure 3A and Figure S6A). With ATP as substrate, we first examined the effect of divalent cations on the hydrolytic activity of GajB. With a 56-nt ssDNA (56-nt ssDNA-F in Table S2) in the reaction, GajB exhibited strong hydrolysis activity in the presence of Mg^2+^ and weak activity in the presence of Mn^2+^ and Ca^2+^, while Zn^2+^, Co^2+^, and Ni^2+^ did not support the ATP hydrolysis activity of GajB (Figure S6B). The optimal metal ion concentration for GajB ATPase activity is 10 mM for Mg^2+^ (Figure S6C) and 2 mM for Mn^2+^ (Figure S6D). The optimal temperature is 30°C (Figure S6E) and the optimal pH is 7 (Figure S6F). GajB hydrolytic activity was inhibited by NaCl or KCl (Figure S7). Therefore, the optimal reaction condition for GajB ATPase activity is established as 20 mM Tris-HCl pH 7, 10 mM MgCl_2_, and 1 mM DTT, at 30°C .

The ATPase activity of GajB is DNA-dependent. By comparison of the effects of various types of DNA or RNA (all at the same mass in each reaction) on the ATPase activity of GajB, including short dsDNA (56 bp, linear), short ssDNA (56 nt, linear), T7 DNA (∼40 kb, linear), pUC19 plasmid DNA, (∼2.7 kb, circular), M13 ssDNA (∼8k nt, circular), PCR-amplified DNA (955 bp, linear), ssRNA (600 nt, linear), and dsRNA (600 bp, linear) (sequences are listed in Table S3), it was clear that GajB is only significantly activated by short dsDNA and ssDNA (Figure 3B), indicating that the ATPase activity of GajB is specifically activated by DNA termini (at the same mass, there are more termini of short DNA than of longer DNA). Further comparison revealed that short ssDNA is more effective than short dsDNA to activate the ATPase activity of GajB (Figure S8A). The stimulation is DNA sequence-independent, as complementary ssDNA showed similar stimulation effect (Figure S8A). To further determine the precise activation signal of GajB, we designed various ssDNAs and compared their effects on GajB ATPase activity. We first found that neither changing the 5’ phosphate to a hydroxyl group nor changing the 3’ hydroxyl group to hydrogen affected the effects of ssDNA on GajB ATPase activity, suggesting that the 5’ phosphate or the 3’ hydroxyl group of DNA is not recognized by GajB (Figure 3C). Attachment of modified groups such as Sp18, idSp, or Cy5 to ssDNA termini also had no obvious effect on GajB activation (Figure S8B). Next, we examined the effects of the length of ssDNA, and found that a 10-nt poly-dT ssDNA is sufficient for GajB activation; further increasing in the DNA length did not further increase the GajB ATPase activity, but shortening to 8-nt and 6-nt gradually decreased the GajB ATPase activity (Figure 3D). Since RNA has no stimulatory effect on GajB (Figure 3B), we added UMPs to the termini of the 10-nt poly-dT to test which side of the DNA termini is crucial for GajB activation. The results showed that the stimulatory effect on GajB ATPase activity gradually decreased with an increasing number of UMPs added at the DNA 3ʹ termini (Figure 3E). In contrast, even a 10-nt poly-U at the DNA 5ʹ termini had no obvious effect on the activation of GajB (Figure 3E). Together, these data revealed that a 10-nt ssDNA with a 3ʹ terminus is the signal for GajB activation.

Among tested NTP and dNTP substrates, GajB showed a high preference for ATP, GTP, dATP, and dGTP to hydrolyze (Figure 3F). To confirm the active site of GajB, mutations of the conserved key residues in the UvrD family, K28A/T29A and D167A/E168A, were constructed and purified (Figure S9A). K28A/T29A or D167A/E168A mutations completely abolished the ATP hydrolysis activity of GajB (Figure S9B), indicating a common mechanism for GajB and UvrD family helicases in ATP hydrolysis. To rule out the effect of the N-terminal His-tag on the activity of GajB, we also expressed the GajB protein with a C-terminal His-tag. The results showed that the GajB protein with a C-terminal His-tag showed almost equal activity to the N-terminal-tagged GajB protein (Figure S9C).

The efficiency of ATP/GTP hydrolysis by GajB was analyzed by TLC. GajB is an efficient ATPase in the presence of ssDNA; 3 µM GajB completely hydrolyzed 4 mM ATP into ADP in 15 min (Figure S10). Similarly, GTP was hydrolyzed by GajB to produce GDP (Figure 3G). We examined the ssDNA in ATP hydrolysis reactions by Native-PAGE and found that the ssDNA was neither stably bound nor processed by GajB under the optimal conditions for ATP hydrolysis (Figure S11).

We also investigated the ATPase activity of the Gabija protein complex by measuring the ATP hydrolysis activity of the following samples: GajA, GajB, *in vitro* assembled GajA/B 1:1 complex (GajA+B), and native GajAB complex (GajA:B∼4:1). The results showed that GajB exhibited the strongest activity and GajA+B and GajAB showed less activity, likely contributed by GajB in the complexes (Figure 3H). The TLC analysis results were similar (Figure S12). The GajA mutations K35A and H320A had no obvious effects on GajAB ATPase activity, while the GajB mutations K28A/T29A and D167A/168A completely abolished the activity of GajAB (Figure 3I).

GajB was initially suspected to function as a helicase based on its homology to UvrD helicases (8). However, in a classic helicase unwinding assay using UvrD as a positive control, GajB displayed no DNA unwinding activity (Figure 3J). Considering GajB may coordinate with GajA and load onto the DNA nicks introduced by GajA, we further employed a nicked dsDNA substrate by annealing three oligonucleotides (Figure S13 and Table S2). However, still no unwinding activity from GajB was detected (Figure S13).

### The interactions between GajA and GajB

Intriguingly, GajA and GajB expressed separately at similar levels (each under the control of the same promoter and RBS, respectively) almost failed in conferring phage resistance (Figure 4A, GajA+B). In the functional native GajAB gene cassette, the expression of GajA was significantly higher than that of GajB (Figure S14). Thus, we speculated that the ratio between GajA and GajB is vital for the function of the Gabija system. To verify the hypothesis, we created new constructs to modify the ratio between Gabija components. In constructs GajAB+A and GajAB+B, extra GajA and GajB were present (expressed under the control of an additional promoter) in the native GajAB cassette, respectively. GajAB+A with additional GajA was as efficient as GajAB in defense against phages. In contrast, GajAB+B with increased amounts of GajB almost lost phage resistance (Figure 4A). The K35A mutation abolished the antiviral defense of the Gabija system (Figure S1B); however, the non-functional K35AGajAB can be rescued by additional WT GajA (Figure 4A). Taken together, these results imply that the GajB level must be lower than the GajA level for Gabija defense. The bacterial growth curves indicated that the Gabija protein has no obvious cytotoxicity to host bacteria in the absence of phage infection (Figure S15).

**Figure 4.**
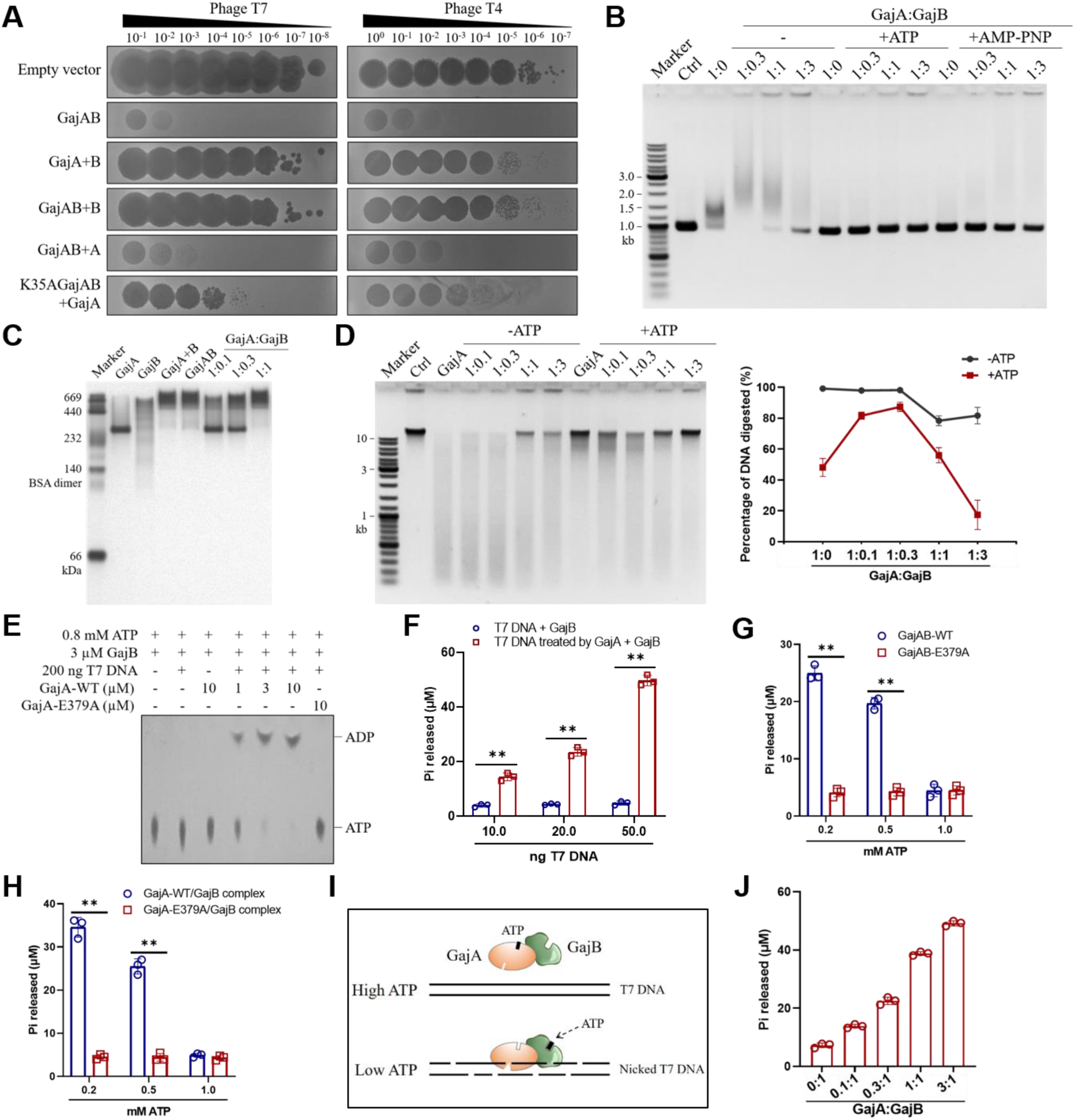
The interactions between GajA and GajB. (**A**) Effects of the relative level of GajA and GajB on phage resistance. Shown are 10-fold serial dilutions of plaque assays with phages T7 and T4 and *E. coli* B carrying various combinations of Gabija components. GajA+B means that the GajA and GajB genes were under the control of two separate promoters. GajAB+B indicates expression of an extra copy of the GajB gene in addition to the native GajAB gene. GajAB+A indicates expression of an extra copy of the GajA gene in addition to the native GajAB gene. K35AGajAB+GajA indicates expression of an extra copy of the GajA gene in addition to the GajAB gene harboring the GajA K35A mutation. Plates were incubated at 37°C overnight before imaging analysis. Images are representative of three replicates. (**B**) Effect of GajB at various GajA/GajB molecular ratios (1:0.3, 1:1, and 1:3 as indicated on the top of the gel) on the pUC19-955 DNA binding of GajA, in the absence or presence of 1 mM ATP or AMP-PNP. Reactions were incubated at 4°C for 30 min. Samples were analyzed via native agarose gel electrophoresis. (**C**) Native-PAGE analysis of purified GajA, GajB, GajA+B, and GajAB, and *in vitro* mixtures of GajA/GajB at various molecular ratios. Each sample containing indicated proteins (45 pmol) was mixed with native gel loading buffer before loading onto the Native-PAGE gel. (**D**) Effect of GajB at various GajA/GajB molecular ratios (1:0.1, 1:0.3, 1:1, and 1:3 as indicated on the top of the gel) on the cleavage of T7 genomic DNA by GajA in the absence or presence of 0.5 mM ATP. DNA digestion was measured using ImageJ software as described in the Materials and Methods. The statistical results are shown on the right. (**E**) TLC analysis of the effect of GajA on the hydrolytic activity of GajB. Reaction mixtures containing 20 mM Tris-HCl pH 8, 2 mM MnCl_2_, and 100 μg/ml BSA in a final volume of 50 µl were incubated at 37°C for 15 min, which were concentrated to 20 μl and used for TLC assays. (**F**) Pi release assay to compare the activation of GajB by intact T7 genomic DNA or T7 genomic DNA nicked by GajA. Reaction mixtures containing 20 mM Tris-HCl pH 8, 2 mM MnCl_2_, 100 μg/ml BSA, 0.2 mM ATP, 1 μM GajB, and 10, 20, or 50 ng purified T7 DNA (with or without 1 μM GajA treated at 37°C for 1 0 min) were incubated at 37°C for 10 min. (**G, H**) Effect of GajA on the hydrolytic activity of GajB in native GajAB complex (**G**) or *in vitro* assembled GajA/B (1:1) complex (**H**) in the presence of various amounts of ATP (0.2, 0.5, and 1 mM). E379A refers to the mutation in the active site of GajA which abolishes its DNA nicking activity. Reaction mixtures containing 20 mM Tris-HCl pH 8, 2 mM MnCl_2_, 100 μg/ml BSA, 100 ng T7 DNA, 1 μM proteins, and 0.2, 0.5, or 1 mM ATP were incubated at 37°C for 15 min. (**I**) Schematic showing the stimulation of GajB ATP hydrolysis by GajA DNA nicking in (**E**–**H**). In the presence of a high concentration of ATP, GajA activity was completely inhibited and GajB hydrolysis could not be activated. In the presence of a low concentration of ATP, GajA exhibited DNA nicking activity and provided DNA termini to activate ATP hydrolysis by GajB. (**J**) Effect of GajA at various GajA/GajB molecular ratios (0:1, 0.1:1, 0.3:1, 1:1, and 3:1) on the ATP hydrolysis activity of GajB. Reaction mixtures containing 20 mM Tris-HCl pH 8, 2 mM MnCl_2_, 100 μg/ml BSA, 100 ng T7 DNA, 1 μM GajB (with various amounts of GajA), and 0.2 mM ATP were incubated at 37°C for 15 min. Pi release was monitored by the PiColorLock™ kit. Data from three independent experiments are presented as the mean ± standard error. Double asterisks (**) indicate *P* < 0.005 as determined by Student’s *t*-test.

We bioinformatically analyzed all 4598 GajAB gene cassettes reported previously (8) and only found bacterial promoters upstream of the GajA gene but not the GajB gene, indicating that GajB shares the same mRNA with GajA (Figure 1A). Nonetheless, we applied RNA-Seq to examine the transcript levels of the GajA and GajB genes (Figure S16). RNA-Seq analyses were performed using *E. coli* B cells expressing the native GajAB gene cassette with or without phage T7 infection. Unexpectedly, the expression level of GajA, either with or without T7 infection, was much higher than that of GajB, based on the counts of reads covering each protein’s coding region (Figure S16A, B). T7 phage infection at a high MOI of 5 slightly reduced the expression level of GajA but significantly increased the expression level of GajB (Figure S16A). In the absence of T7 infection, the expression level of GajA was over 20 times higher than that of GajB. In the presence of a high MOI T7 infection, the expression level of GajA was several times higher than that of GajB (Figure S16A). We speculate that this regulation may be caused by the early termination of transcription in the GajB gene’s early region. Indeed, bacterial transcription terminator-like structures were predicted in the early region of the GajB gene downstream of the GajA stop codon using the RNAfold WebServer (Figure S16C). To experimentally support this hypothesis, we expressed the same GajAB gene cassette under the control of either a bacterial promoter (T5) or a phage promoter (T7) in *E. coli* BL21(DE3). In the former situation, the *E. coli* polymerase performing the transcription of the GajAB gene cassette should recognize the potential terminator in the GajB gene, while in the latter situation, the phage T7 RNA polymerase may not be efficiently terminated. Consistent with this hypothesis, expression using *E. coli* RNA polymerase resulted in a much higher level of GajA relative to GajB, whereas expression using T7 polymerase produced GajA and GajB at similar levels (Figure S16D).

Unlike GajA, GajB exhibited no significant binding to the pUC19-955 dsDNA containing GajA recognition sites (Figure S17A), even in the presence of ATP or AMP-PNP (Figure S17B). The binding to the 56-nt ssDNA and 56-bp dsDNA that trigger the ATPase activity by GajB was also not detectable by EMSA (Figure S17C). Since GajB forms a complex with GajA, we examined the effects of GajB on the DNA binding by GajA. The results showed that at a molecular ratio of GajA/GajB = 1:0.3, GajB enhanced the DNA binding of GajA (Figure 4B); however, when the level of GajB was equal to or exceeded that of GajA, GajB inhibited the DNA binding of GajA (Figure 4B). At low molecular ratios, GajB not only enhanced the DNA binding of GajA but also retarded the gel shift of bound DNA, indicating that GajB might be incorporated into the GajA/DNA complex (Figure 4B). ATP or non-hydrolyzable AMP-PNP at 1 mM completely inhibited the DNA binding of GajA, even in the presence of GajB (Figure 4B), implying that the Gabija system was suppressed under physiological NTP concentrations (Figure S15). Further, we compared the DNA binding of the GajA and GajAB native complex in the presence of ATP. We found that at low ATP concentrations (0.25 and 0.5 mM), GajAB bound DNA more strongerly than GajA alone (Figure S17D), indicating that GajB in the complex enhanced the DNA binding of GajA and attenuated the inhibition from ATP. The K35A and H320A mutations in the GajAB complex also attenuated the inhibitory effects of ATP on DNA binding (Figure S17E–G).

GajA with an N-terminal 6×His-tag has a molecular weight of about 69 kDa. In Native-PAGE, GajA runs as an evident band corresponding to about 276 kDa, indicating that GajA forms a multimer (likely a tetramer) (Figure 4C). By contrast, GajB tends to form polymers of various and gradually increasing molecular weight (Figure 4C). When mixed, GajB interacts with GajA to form large complexes, as shown near the top of the gel (Figure 4C).

Subsequently, we examined the effects of GajB on the DNA cleavage activity of GajA. The robust DNA nicking activity of GajA results in the damage of T7 genomic DNA (Figure 4D). In the absence of ATP, the addition of GajB at various molecular ratios inhibited the cleavage activity of GajA (Figure 4D). However, in the presence of 0.5 mM ATP, GajB supplied at a molecular ratio to GajA lower than 1:1 significantly stimulated the cleavage activity of GajA (Figure 4D), implying the key role of GajB in assisting GajA activation in the presence of cellular NTP. Intriguingly, activation of GajB ATP hydrolysis in the complex by adding extra ssDNA did not further stimulate DNA cleavage by GajA in the presence of ATP (Figure S18), possibly due to the inhibitory effects of the excess ssDNA on GajA activity. However, active site mutations in GajB reduced the DNA cleavage by the GajAB complex in the presence of ATP, indicating that ATP hydrolysis by GajB contributed to GajA activation (Figure S19), but in this assay no additional DNA termini were supplied to activate the ATP hydrolysis of GajB.

We initially hypothesized that GajA was the antiviral effector and GajB the regulator. However, the above results that ATP hydrolysis of GajB was activated in the GajAB complex without additional DNA termini led us to hypothesize that the regulation may also take place in an opposite way, in which GajA is the regulator to provide DNA termini to activate GajB, while GajB acts as an antiviral effector by nucleotide depletion. The DNA nicking activity of GajA makes it an ideal provider of DNA termini, and nucleotide depletion is known to mediate defense against phages (52–56). Therefore, we first examined the effects of GajA DNA nicking activity on the ATP hydrolysis activity of GajB through TLC analysis (Figure 4E). GajB alone exhibited no obvious ATP hydrolysis activity, even in the presence of T7 genomic DNA. Adding GajA without DNA also failed to activate GajB. However, the presence of both GajA and T7 genomic DNA effectively activated the ATP hydrolysis activity of GajB, and the activation effect was more obvious with increasing levels of GajA; the mutation E379A disrupting the DNA nicking activity of GajA abolished the activating effects of GajA on GajB (Figure 4E). The Pi release assay showed that intact T7 genomic DNA was not able to stimulate the ATPase activity of GajB. However, equal amounts of T7 genomic DNA treated by GajA significantly activated GajB to hydrolyze ATP (Figure 4F). These assays clearly show that the DNA termini produced on T7 genomic DNA by GajA nicking activity stimulated the ATP hydrolysis activity of GajB. Then, we tested the ATPase activity of GajB in the GajAB form. The results showed that GajAB-WT exhibited much higher activity than GajAB-E379A (the mutation E379A was in the active site of GajA to inactivate its DNA nicking activity) in the presence of T7 genomic DNA and ATP at low concentration, while they display no obvious difference in the presence of 1 mM ATP (Figure 4G). We also examined the *in vitro* assembled GajA/GajB complex and obtained similar results: At low ATP concentrations, the DNA nicking by GajA significantly stimulated ATP hydrolysis by GajB, while the mutation inactivating GajA abolished the stimulatory effects (Figure 4H). If the initial ATP concentration was too high to release GajA activity, GajB activation was abolished (Figure 4G, H). Graphical models (Figure 4I) demonstrated the regulatory effects of GajA on GajB. In the presence of high concentrations of ATP, GajA activity was completely inhibited, and the GajB hydrolysis activity could not be activated effectively. At lower concentrations of ATP, GajA exhibited DNA nicking activity, and produced sufficient DNA termini, which effectively activated GajB ATP hydrolysis (Figure 4I). Unlike the inhibitory effects of GajB on GajA activity at a molecular ratio of 1:1, GajA stimulated the activity of GajB more efficiently at a molecular ratio of 1:1 or higher (Figure 4J). Taken together, at a GajA/GajB molecular ratio of >1:1, GajA and GajB mutually enhance each other’s activity in the presence of DNA and ATP, resulting in both DNA cleavage and ATP hydrolysis.

### Effects of Gabija defense on the cellular level of DNA and NTP

qPCR analysis was conducted to track the variation of phage and host DNA at various time points following phage T7 infection. The results showed that T7 DNA levels are significantly lower in Gabija-containing cells compared to Gabija-lacking cells (∼8.5-fold reduction at 10 min post-infection and ∼20-fold reduction at 20 min post-infection) (Figure 5A). Meanwhile, the accumulation of host DNA was also decreased in Gabija-containing cells (Figure 5A). The integrity of total genomic DNA at various time points following phage T7 infection was visualized on the gel. The results showed that both the host and phage genomic DNA were damaged by the Gabija defense system, based on the comparison of the DNA integrity on the gel in the absence and presence of the Gabija system after a 20-min phage infection (Figure S20). Furthermore, the effects of T7 infection and the Gabija defense system on bacterial cells were visualized using fluorescence microscopy. Compared with cells lacking the Gabija system, cellular DNA in a large portion of cells carrying the Gabija system was either partially or completely degraded at 20 min post-infection; the degradation was more obvious at 30 min post-infection (Figure 5B and Figure S21). These *in vitro* and *in vivo* results are consistent with our previous study, showing that GajA catalyzes DNA nicking on both T7 and host genomic DNA *in vitro* (45) and the abortive infection mechanism.

**Figure 5.**
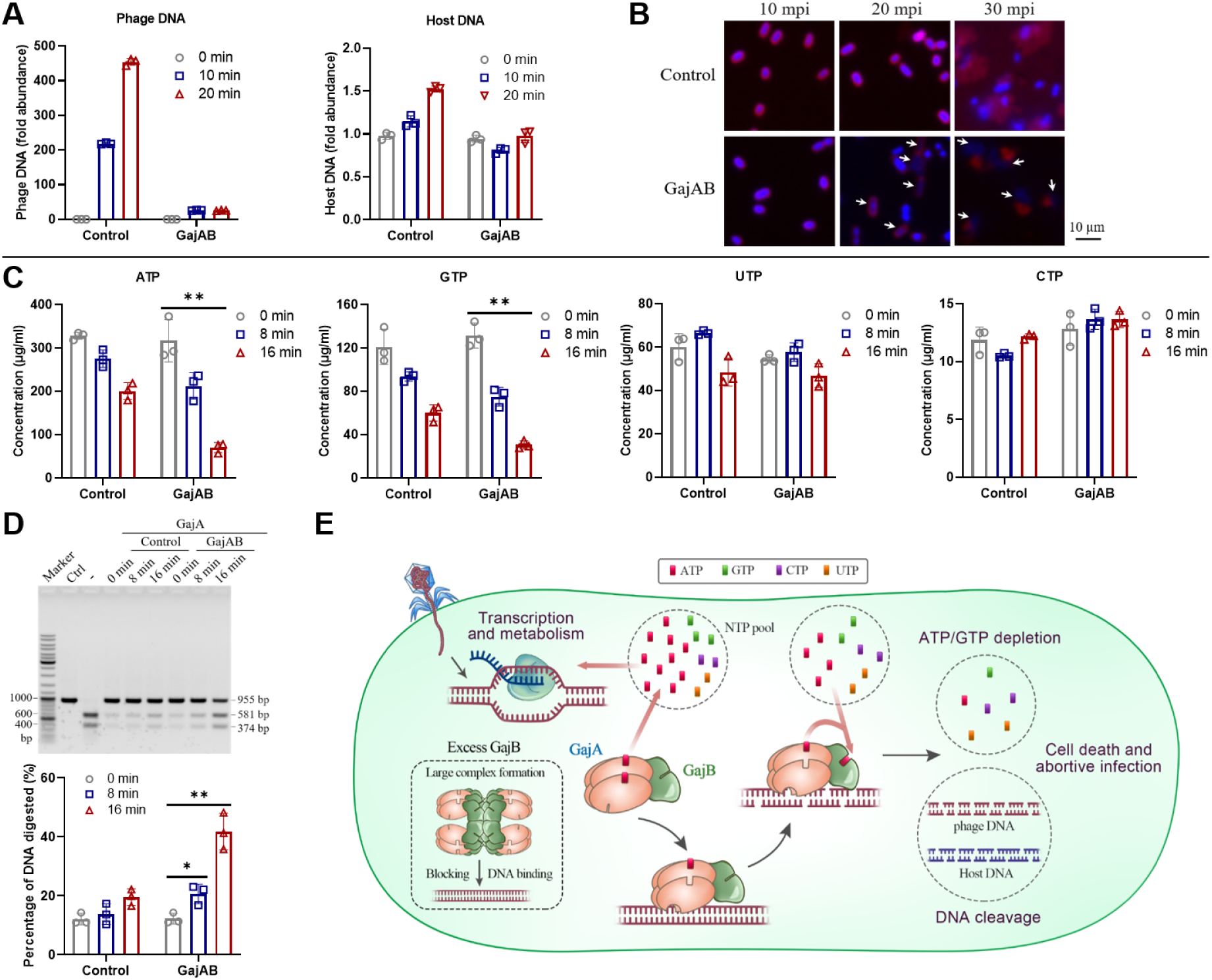
Antiviral mechanism of the Gabija defense system. (**A**) Relative abundance of phage DNA and host DNA at various time points following phage infection as measured by qPCR. Relative DNA abundance at each time point was normalized against the control (empty vector) at 0 min post-infection. The differences in *Ct* between the control (empty vector, 0 min) and the samples (Δ*Ct*) were calculated, and 2^(−Δ^*^Ct^*^)^ for each sample is shown. (**B**) Fluorescence microscopy of *E. coli* B cells harboring the GajAB gene cassette or empty vector (control). DAPI staining (DNA) is shown in blue, and FM4-64 staining (membranes) is shown in magenta. White arrows indicate the degradation of host DNA. Separate channels are shown in Figure S21. Scale bar: 10 µm. (**C**) Concentrations of NTP in cell lysates extracted from T7-infected cells with or without the Gabija system, as measured by UHPLC-MS/MS with synthesized standards. Cells were infected by phage T7 at an MOI of 2. (**D**) Effects of T7-infected cell lysates (with or without the Gabija system) on GajA cleavage activity. Cell lysates were prepared as described in the Materials and Methods section. pUC19*-*955 DNA (125 ng) was incubated with 0.2 µM GajA in a final volume of 10 µl in the optimal reaction buffer at 37°C for 5 min with or without various cell lysates. Ctrl refers to the control without protein. “-” indicates the reaction without cell lysates. DNA digestion was measured using ImageJ software as described in the Materials and Methods section, and the statistical analysis results are shown on the right. Bar graphs represent the mean ± SD of three independent experiments. Single (*) and double (**) asterisks indicate *P* < 0.05 and *P* < 0.005, respectively, as determined by Student’s *t*-test. (**E**) Model for the Gabija antiviral defense mechanism. During phage invasion, phage transcription and metabolism reduce cellular A/GTP levels, which relieves the inhibitory effects on the DNA binding of GajA. Once the DNA nicking of both phage and host DNA by GajA occurs, it provides DNA termini to the complexed GajB to activate its (d)ATP/(d)GTP hydrolysis activity. By maintaining a specific molecular ratio in the complex, GajB further enhances the DNA cleavage by the complexed GajA through A/GTP hydrolysis and allosteric effects. The mutual activation circle between GajA and GajB results in a cascade suicide effect of DNA damage and A/GTP depletion to abort phage infection.

We next detected the effects of Gabija defense on the cellular NTP pool size using UHPLC-MS/MS. After phage T7 infection, the concentrations of ATP and GTP were significantly decreased in cells with or without the Gabija system (Figure 5C), likely due to the consumption of these two pivotal nucleotides in many metabolic processes in addition to transcription. With T7 infection, the presence of the Gabija system further reduced the ATP and GTP concentrations (Figure 5C). The Gabija defense system against T7 infection finally reduced the ATP concentration to ∼0.67 mM (a 78% reduction compared to normal cellular levels) and the GTP concentration to ∼0.28 mM (a 77% reduction compared to normal cellular levels) at 16 min post-infection. However, UTP and CTP levels were not affected by the Gabija defense system (Figure 5C), consistent with the substrate specificity of GajB. dNTPs with much lower concentrations than cellular NTPs (57) are less significant to affect GajA and were not investigated in this study. ATP and GTP are the most abundant cellular NTPs (above 5.2 mM) (57). Reduction of their levels in cells after phage infection will initially activate GajA, which will be further enhanced by the A/GTP hydrolysis activity of GajB. To further confirm GajA activation after phage infection, we examined the effects of cell lysates collected at various time points following phage infection on GajA cleavage activity.

GajA exhibited stronger DNA cleavage activity in cell lysates collected after phage T7 infection, especially in cell lysates expressing GajAB (Figure 5D), implying GajA is activated during Gabija defense *in vivo*.

## DISCUSSION

### GajA and GajB are both signal sensors and antiviral effectors for Gabija defense

In our previous study, we revealed that GajA is a site-specific DNA nicking enzyme and is negatively regulated by high nucleotide concentrations (45). The physiological concentration of ATP is over 3 mM and the total NTP concentration is above 6.2 mM in *E. coli* at the mid-log phase (57), while GajA nuclease activity is seriously inhibited by 1 mM ATP *in vitro* (Figure 2A). Therefore, the robust DNA nicking activity of GajA should be suppressed by NTP at physiological concentrations. Indeed, overexpression of Gabija proteins results in no toxicity to *E. coli* (Figure S15), indicating that GajA activity is tightly suppressed *in vivo*. Our results demonstrated that ATP inhibits GajA through its specific binding to the DNA recognition site (Figure 2). The sensing of NTP concentration of GajA is mediated by its ATPase-like domain, as mutations of K35 and H320 in this domain decreased the inhibitory effects of ATP on DNA binding (Figure S17E). Sensing of nucleotide depletion is a novel and straightforward strategy, especially to virulent phages such as T7 and T4. Indeed, a drastic reduction in the levels of ATP and GTP, two of the most abundant cellular NTPs (above 5.2 mM combined) was observed after T7 infection (Figure 5C).

However, cellular NTP levels are not completely depleted by phage transcription and metabolism (Figure 5C), thus GajA activation requires assistance. The DNA terminus-dependent A/GTPase GajB (Figure 3B–E) complexed with GajA may conveniently sense the DNA termini provided by GajA to further hydrolyze A/GTP (Figure 4E–H). The depletion of A/GTP by both phage metabolism and GajB hydrolysis activates GajA to degrade phage and host DNA (Figure 5D). Although the partial degradation of phage and host DNA by GajA (Figure 5A) cannot account for the robust antiviral effect of the Gabija system, the combination of DNA degradation by GajA and A/GTP depletion by GajB could result in efficient cell death.

Although the DNA termini produced by GajA serve as the main signal to activate GajB, we cannot exclude the possibility that the DNA termini produced in the robust phage DNA metabolism (58, 59) serve as an initial signal to activate GajB, just like the initial signal of NTP consumption by phage transcription and metabolism to activate GajA. Once both GajA and GajB in the Gabija complex are initially activated, they may mutually activate each other. This mutual activation results in a cascade effect, in which DNA cleavage and A/GTP depletion are both signals and effects, while GajA and GajB are both signal sensors and antiviral effectors, distinguishing the Gabija system from known defense systems.

GajA alone forms uniform multimers, while GajB alone forms polymers (Figure 4C). When mixed at a molecular ratio of 1:1, they together form high-molecular-weight large complexes (Figure 4C), which decreases the DNA binding (Figure 4B). These results indicate that there are multiple (at least two) protein binding sites in GajB, and these sites can bind to either GajA or another GajB molecule. Based on these results, we proposed a mechanism to explain why excess GajB leads to the loss of antiviral function: In the case of the native GajA/GajB complex, the binding sites of GajB are fully occupied by GajA within individual complexes; while in the GajA/GajB complex containing more GajB than GajA, empty binding sites of GajB are exposed and can bind to other complexes, which could result in the formation of large complexes and block the DNA binding and cleavage by GajA. It is likely that the lower expression level of GajB, relative to GajA, is achieved through the regulation of transcription termination, given that GajB shares the same mRNA as GajA (Figure S16).

Collectively, our data suggest a model for the antiviral mechanism of the Gabija immune system (Figure 5E). The specific DNA binding of the GajA endonuclease is fully inhibited by nucleotides during the physiological state. After phage invasion, the robust phage transcription and metabolism consume cellular NTP. The reduction in NTP concentration is sensed by the ATPase-like domain of GajA as an initial signal, which allosterically activates the TOPRIM domain of GajA to bind to and cleave both the phage and host DNA. Meanwhile, the DNA termini produced by the DNA nicking of GajA and perhaps also from the DNA metabolism of invading phages are sensed by GajB and activate its (d)A/(d)GTP hydrolysis activity. The activated GajA and GajB function as a complex, in which GajA and GajB are both signal sensors and antiviral effectors to enhance each other’s activity. The resulting cascade effect of both DNA damage and nucleotide depletion leads to an efficient abortive infection defense against virulent bacteriophages. Since GajB possesses multiple protein interaction sites, the molecular ratio between GajB and GajA must remain low to prevent the formation of large and non-functional complexes which block DNA binding.

### Two collinear GajB proteins expressed by the *B. cereus* VD045 GajAB gene cassette

By careful dissection of the products of the native *B. cereus* VD045 GajAB gene cassette in *E. coli*, we unexpectedly found that the purified proteins contained GajA and two forms of GajB of close sizes (Figure S22A). Protein N-terminal sequencing revealed that the shorter GajB (hereafter referred to as GajB-S) was translated from the previously predicted start codon (8), while the larger GajB (referred to as GajB in this work) was translated from another start codon located in the GajA gene (Figure S22B). GajB and GajB-S share the same coding and amino acid sequences except that GajB possesses five additional amino acids (MIEDE) in its N-terminus (Figure S22C, D). Thus, the Gabija system encoded by the *B. cereus* VD045 gene cassette contains three protein components: a GajA endonuclease and two collinear GajB proteins (GajB and GajB-S) (Figure S22B). Mutation of the start codon of either GajB or GajB-S specifically eliminates its expression (Figure S22E). The mutations in GajA and GajB investigated in this work did not change the expression of the GajAB gene cassette (Figure S22E).

If GajB was removed by mutating its initiation codon GTG to GCG, the combination of co-expressed GajA and GajB-S (GajAB-GCG) almost lost its antiviral function (Figure S23A). In contrast, the removal of GajB-S by mutating its initiation codon ATG to ATC from the native Gabija complex (GajAB-ATC) had no obvious effect on its antiviral function (Figure S23A). Therefore, GajB-S is functionally redundant, at least for the antiviral mechanism investigated in this work. Consistent with its redundancy in phage resistance, GajB-S showed only minimal NTP hydrolysis activity (Figure S23C–E).

Nonetheless, we cannot exclude the possibility that the two collinear GajBs may play a regulatory role in certain situations or in the Gabija defense systems of other bacteria. Therefore, we bioinformatically analyzed the potential collinear GajBs in all predicted Gabija gene cassettes (8). First, bioinformatics analysis did not reveal any potential promoters in front of the *GajB* gene in all 4598 GajAB gene cassettes reported previously (8), indicating that GajB (and GajB-S) shares the same mRNA with GajA and is translated under the control of its own ribosome-binding site (RBS) and start codon. Further, among the 4598 predicted Gabija systems (8), the bioinformatics analysis predicted that 970 (21.1%) systems may code for two collinear GajBs under the control of individual but proximal RBSs and start codons (Figure S24; detailed in Table S4). Among these 970 Gabija operons, we experimentally investigated the protein expression of two operons from *Burkholderia pseudomallei* 668 (8, 60) and *Bacillus cereus* HuB5-5 (8). However, either operon only expressed a single GajB in *E. coli* (Figure S25). We could not exclude the possibility that their expression may be differentially regulated in native hosts.

## SUPPLEMENTARY DATA

Supplementary data are available online.

## DATA AVAILABILITY

Data that support the findings of this study are available within the main text and the supplementary data. All data are available from the corresponding author upon request.

## ACKNOWLEDGEMENTS

We thank all lab members for helpful discussion. This work is supported by the National Natural Science Foundation of China (grant 32150009 to B.Z., 32100025 to R.C.) and Fund from Science, Technology and Innovation Commission of Shenzhen Municipality (grant JCYJ20210324115811032 to B.Z.). Funding for open access charge: National Natural Science Foundation of China.

## AUTHOR CONTRIBUTION STATEMENT

R.C. and B.Z. conceived the project and designed the experiments. R.C. carried out the experiments. R.C., F.T.H., X.L.L., Y.Y., B.B.Y., X.L.W. and B.Z. analyzed the data. R.C. and B.Z. wrote the manuscript. All authors discussed the results and contributed to the final manuscript.

## Figure legends

## MATERIALS AND METHODS

### Materials

Oligonucleotides, primers, and genes were synthesized by Genscript Corporation (Nanjing, China). The Gibson assembly kit and RNA purification kit were purchased from New England BioLabs. PrimeSTAR Max DNA Polymerase was obtained from TaKaRa. The DNA purification kit was obtained from Axygen. Ni-NTA resin was obtained from Qiagen. Preparative Superdex S200 (catalog no. 17-1043-01) for gel filtration was obtained from GE Healthcare. ATPase activity was measured with the PiColorLock™ phosphate detection system kit (Expedeon). DNA ladder (#SM0331) and protein ladder (#26619) were obtained from Thermo Scientific™. The protein marker used for Native-PAGE was obtained from Real-Times Biotechnology (Beijing, China).

### Cloning, expression, and purification of Gabija proteins

The following predicted coding sequences were cloned into the pET28a vector harboring an N-terminal 6×His -tag using Gibson Assembly Cloning Technology: GajA (GenBank accession number: MW659467, residues 1–578), GajB-S (GenBank accession number: MW659468, residues 1–494), GajB (GenBank accession number: OM891105, residues 1–499), GajAB (located at 94190–97412 of the *B. cereus* VD045 genome [AHET01000033]), GajA+B-S (GajA and GajB-S genes under the control of separate promoters), and GajA+B (GajA and GajB genes under the control of separate promoters) (61). The constructs were transformed into *E. coli* BL21(DE3) cells, which were cultured in 1 L LB medium containing 50 μg/ml kanamycin at 37°C for ∼3 h to an OD_600_ of 0.6–0.8, and then induced with 0.2 mM IPTG for 20 h at 12°C.

The cells were harvested, and resuspended in lysis buffer (20 mM Tris-HCl pH 7.5, 300 mM NaCl, and 0.3 mM DTT), and lysed by ultrasonication. Supernatant was collected after centrifugation (13,000 rpm, 4°C, 1 h), filtered with a 0.45-μm filter, and loaded onto a Ni-NTA agarose column pre-equilibrated with 10 volumes of elution buffer (20 mM Tris-HCl pH 7.5 and 300 mM NaCl), and then the column was washed with 10 volumes of elution buffer containing 20 mM and 50 mM imidazole, respectively. The majority of GajA was eluted off the column by elution buffer containing 120 mM imidazole. Collected elutes were concentrated to 2.8 ml by a Millipore Amicon Ultra-15 (30,000 MWCO) and further purified by molecular sieve chromatography on a preparative Superdex S200 column. Fractions containing pure Gabija proteins were concentrated again. Finally, Gabija proteins were dialyzed against a storage buffer (50 mM Tris-HCl pH 7.5, 100 mM NaCl, 1 mM DTT, 0.1 mM EDTA, 50% glycerol, and 0.1% Triton X-100).

Active site mutations were introduced using the Gibson assembly method (61) (primers for cloning are listed in Table S5), and mutants were expressed and purified using the same procedure as detailed above.

### Protein analysis

The concentrations of purified proteins were determined by a Bradford protein quantitative kit (Bio-Rad), and protein purity was analyzed by 10% SDS-PAGE followed by staining with Coomassie blue (Bio-Rad), with BSA as a standard. In addition, Native-PAGE analysis was performed using 8% gels. The protein marker used for Native-PAGE was purchased from Real-Times Biotechnology (Beijing, China). Proteins were analyzed by staining the gels with Coomassie blue (Bio-Rad).

### Measurement of bacterial growth curve

The sequences of the genes were cloned into the pQE82L vector. The recombinant vectors were transformed into *E. coli* B (ATCC^®^ 11303™). Empty vector was used as a control. A single bacterial colony was picked from a fresh LB agar plate and grown in LB broth supplemented with ampicillin (100 µg/ml) at 37°C . Overnight cultures were diluted 1:100 into liquid LB medium containing ampicillin (100 µg/ml) and grown at 37°C for 2 h to an OD_600_ of about 0.6. Subsequently, the cultures were divided into two portions supplemented with or without 1 mM IPTG. Every 1 h, aliquots from the cultures were taken to monitor the OD_600_ using a NanoPhotometer^®^ (Implen). Graphs were produced using SigmaPlot software.

Phage infection time-course assays were performed in liquid media. *E. coli* B cells harboring the GajAB gene cassette or the empty vector were grown at 37°C with vigorous shaking and infected with phages T7 or T4 at *t* = 0 at a multiplicity of infection (MOI) of 0, 0.05, or 5 in three replicates. Optical density at 600 nm (OD_600_) was measured every 10 min for a period of 600 min (for phage T7) or 360 min (for phage T4).

### Plaque assays

Phages were propagated by picking a single phage plaque into a liquid culture of *E. coli* B grown at 37°C to an OD_600_ of ∼0.4 in LB medium until culture collapse. The culture was then centrifuged for 4 min at 12,000 rpm and the supernatant was filtered through a 0.2-µm filter to remove remaining bacteria and bacterial debris. Lysate titer was determined using the small drop plaque assay method as previously described (62, 63).

Plaque assays were performed as previously described (8,62). The sequences of the genes were cloned into the pQE82L vector. The recombinant vectors were transformed into *E. coli* B. A single bacterial colony was picked from a fresh LB agar plate and grown in LB broth containing ampicillin (100 µg/ml) at 37°C to an OD_600_ of ∼0.4. Protein expression was induced by the addition of 0.2 mM IPTG. After further growth for ∼1 h, 500 μl of the bacterial cultures was mixed with 14.5 ml of 0.5% LB top agar, and the entire samples were poured onto LB plates containing ampicillin (100 µg/ml) and IPTG (0.1 mM). Plates were spotted with 4 μl of the three phages diluted in LB at eight 10-fold dilutions, namely, 10^−1^–10^−8^ for T7 and 10^0^–10^−7^ for T4 and T5. Plates were incubated at 37°C overnight and then imaged.

### Preparation of double-stranded DNA

The oligonucleotides used in this study were synthesized by Genscript Company. The synthetic double-stranded DNA (dsDNA) was prepared by mixing equimolar amounts (20 µM) of complementary single-stranded oligonucleotides in a total volume of 100 µl in annealing buffer (10 mM Tris-HCl pH 7.4 and 50 mM NaCl). Complementary oligonucleotides were annealed by heating at 95°C for 5 min followed by gradient cooling to 25°C over a period of 110 min.

### Hydrolytic activity assays

Hydrolytic activity was determined by the PiColorLock™ phosphate detection system kit (Expedeon), which measures the amount of free phosphate released. The reactions were performed in hydrolysis reaction buffer (20 mM Tris-HCl pH 7.0, 10 mM MgCl_2_, and 1 mM DTT) with 0.5 mM ATP, 0.5 µM ssDNA (Table S2), and 0.5 µM protein at 30°C for 15 min, unless stated otherwise. The reaction was stopped by adding the malachite green solution at a sample:dye ratio of 4:1. Subsequent processing was performed following the kit manual and the samples were quantified by a NanoPhotometer^®^ (Implen) at 650 nm.

### Thin layer chromatography

Hydrolytic activity of the reaction products was detected by thin layer chromatography (TLC). Unless mentioned otherwise, the reaction was performed in optimal reaction buffer with 4 mM ATP, 3 µM ssDNA, and 3 µM protein at 30°C for 15 min. Aliquots (1 μl) of samples were spotted onto a polyethyleneimine cellulose TLC plate (Merck, Germany) and developed with a solution containing 1 M formic acid and 0.8 M LiCl as previously described (64). ATP, ADP, and AMP were used as the standards for TLC analysis.

### Helicase activity assays

Unless mentioned otherwise, GajB or GajB-S was incubated at 30°C with the helicase substrates (50 nM oligonucleotides) in 10 μl of the reaction buffer (20 mM Tris-HCl pH 7, 10 mM MgCl_2_, and 1 mM DTT) supplemented with 1 mM ATP. After a 10-min incubation, reactions were stopped by adding 2 µl of 6× loading dye containing 20 mM EDTA. Samples were analyzed by 10% Native-PAGE in Tris-Borate-EDTA (TBE) buffer. All oligonucleotides used in this study are listed in Table S2. The gels were stained with ethidium bromide and imaged on a Typhoon TRIO+ variable mode imager (GE Healthcare).

### Electrophoretic mobility shift assay

Binding assays were carried out with pUC19-955 DNA or oligonucleotides in binding buffer (20 mM Tris pH 8.0 and 5 mM CaCl_2_ or MgCl_2_). A certain amount of DNA was allowed to bind with various concentrations of GajA (0.3, 1, and 3 μM). Protein and DNA were mixed and incubated at 30°C or 4°C for 30 min. Reactions were stopped by adding 2 µl of 6× loading dye. Samples were analyzed by native agarose gel electrophoresis.

### RNA-Seq analysis

*E. coli* B mid-log cells harboring the GajAB gene cassette were infected with phage T7 at an MOI of 0, 0.05, or 5, and samples were taken at 20 min post-infection (each sample with three replicates). RNA was extracted from three biological replicates using the TRIzol^®^ reagent according to the manufacturer’s instructions (Invitrogen), and genomic DNA was removed using DNase I (NEB). The RNA-seq transcriptome libraries were constructed using the TruSeq^TM^ RNA sample preparation kit (Illumina, San Diego, CA, USA) with a total of 2 µg RNA. Ribosomal RNA (rRNA) depletion was performed using the Ribo-Zero Magnetic kit (epicenter), and then all mRNAs were broken into short (200-nt) fragments by adding fragmentation buffer. High-throughput sequencing was performed using an Illumina HiSeq×TEN platform (2 × 150 bp read length). The raw reads in FASTQ format were trimmed and quality controlled using the Illumina GA Pipeline (version 1.6), in which 150-bp paired-end reads were obtained. A Perl program was written to select clean reads by removing low-quality sequences. Clean reads (>4 Gbp for each sample) with Q30 > 93% were then aligned to the genome of *E. coli* K12 (NC_000913.3) and plasmid sequences using Bowtie2 (http://bowtie-bio.sourceforge.net/bowtie2/index.shtml) after removing the reads from phage T7 (NC_001604.1). The gene expression levels were calculated using the fragments per kilobase of transcript per million mapped reads (FPKM) method. The accession number of the raw RNA-Seq reads generated in this study is PRJNA96189X.

### DNA cleavage assays

To investigate the synergistic effects of GajA and GajB, DNA cleavage experiments were performed in 10-μl reaction volumes with 200 ng of phage T7 DNA and 0.3 μM proteins in reaction buffer (20 mM Tris-HCl pH 8.0, 5 mM MgCl_2_, and 0.1 mg*/*ml BSA, with optional addition of 0.5 mM ATP). The proportions of GajA and GajB were 1:0.1, 1:0.3, 1:1, and 1:3, and compared with GajA alone. Reactions were carried out at 37°C for 10 min and then stopped by adding 2 µl of 6× loading dye. Samples were analyzed by native agarose gel electrophoresis. After ethidium bromide staining, the signal of the initial DNA substrate was determined and quantified using ImageJ software (65). To determine the ratio of DNA degradation, the intensity of the intact DNA substrate band in each lane was compared to the intensity of the intact DNA band in the protein-free control lane.

### Quantitative real-time PCR

*E. coli* B mid-log cells harboring the GajAB gene cassette or the empty vector were infected with phage T7 at an MOI of 0.5, and samples were taken at 0, 10, and 20 min post-infection and spiked with an equal volume of *Bacillus subtilis* (ATCC^®^ 6051™). Samples were heated at 95°C for 10 min immediately after sampling to lyse the cells and release the DNA. Then samples were diluted 1:10 in sterile water, and equal volumes of which were used for quantitative PCR (qPCR). qPCR analysis was performed using ChamQ SYBR qPCR Master Mix (Q311, Vazyme) with a CFX Connect^TM^ Real-Time System (Bio-Rad). *Bacillus subtilis* (ATCC^®^ 6051™) was used as a spike-in control. DNA abundance was normalized to the control at 0 min post-infection. The differences in the *Ct* between the normalizer and the samples (Δ*Ct*) were calculated and 2^(−Δ^*^Ct^*^)^ for each sample is shown. Triplicate measurements were taken for each of at least two independent trials. The sequences of phage-specific primers and host-specific primers are listed in Table S2.

### Fluorescence microscopy

Fluorescence microscopy assays were performed as previously described (29, 66). Briefly, overnight cultures of *E. coli* B harboring the GajAB gene cassette or empty vector were diluted 1:100 and grown at 37°C until reaching an OD _600_ of 0.3–0.4. The cultures were infected with phage T7 at an MOI of 2.

Samples of each culture were taken at 10, 20, and 30 min post-infection and stained with 4’,6-diamidino-2-phenylindole (DAPI; 4 µg/mL) and FM4 -64 (2 µg/ml) (67) for imaging with fluorescence microscopy.

### UHPLC-MS/MS quantification of nucleotides

Overnight cultures of *E. coli* B harboring the GajAB gene cassette or empty vector were diluted 1:100 in 200 ml LB medium and grown at 37°C until reaching an OD _600_ of 0.3–0.4. The cultures were infected with phage T7 at a final MOI of 2. Next, samples (50 ml) were taken at 0, 8, and 16 min post-infection and centrifuged for 5 min at 7,200 *g*. Pellets were flash frozen using liquid nitrogen. The pellets were resuspended in 500 µl of cold extraction buffer (acetonitrile, methanol, water, formic acid = 2:2:1:0.02, v/v/v/v) containing 100 ng/ml AMP-13C5. The samples were further treated for five times with 1 min ultrasonication in ice-water and 1 min at −80°C . The homogeneous mixture was centrifuged at 15,000 *g* for 15 min at 4°C, and the supernatant was collected, and the dried residue was reconstituted in 100 μl of 50% acetonitrile prior to performing UHPLC-MS/MS analysis. UHPLC-MS/MS analysis was performed on an Agilent 1290 Infinity II UHPLC system coupled to a 6470A Triple Quadrupole mass spectrometry (Santa Clara, CA, United States). Samples were injected onto a SunArmor NH2 column (150 mm × 2.1 mm, 3 μm, ChromaNik Technologies Inc.) at a flow rate of 0.3 ml/min. The eluted analytes were ionized by electrospray ionization source in positive mode (ESI+). Multiple reaction monitoring (MRM) was applied to determine the contents of nucleotides and internal standards. Raw data were processed with MassHunter Workstation Software (version B.08.00, Agilent) using the default parameters and manual inspection to ensure the qualitative and quantitative accuracies of each nucleotide.

### Cell lysate preparation

Overnight cultures of *E. coli* B harboring the GajAB gene cassette or empty vector were diluted 1:100 in 200 ml LB medium and grown at 37°C until reaching an OD _600_ of 0.3–0.4. The cultures were infected with phage T7 at a final MOI of 2, and 15-ml samples were taken at 0, 8, or 16 min post-infection. The pellets were re-suspended in 200 µl of water, lysed by ultrasonication, and then centrifuged at 13,000 *g* for 15 min at 4°C . The supernatant was transferred to an Amicon Ultra-0.5 Centrifugal Filter Unit 3 kDa (Merck Millipore, Catalogue no. UFC500396) and centrifuged at 10,000 *g* for 20 min at 4°C. The filtrate was taken and used for the subsequent experiments.

